# Enviromic-based Kernels Optimize Resource Allocation with Multi-trait Multi-environment Genomic Prediction for Tropical Maize

**DOI:** 10.1101/2021.06.11.448049

**Authors:** Raysa Gevartosky, Humberto Fanelli Carvalho, Germano Costa-Neto, Osval A. Montesinos-López, José Crossa, Roberto Fritsche-Neto

## Abstract

Genomic prediction (GP) success is directly dependent on establishing a training population, where incorporating envirotyping data and correlated traits may increase the GP accuracy. Therefore, we aimed to design optimized training sets for multi-trait for multi-environment trials (MTMET). For that, we evaluated the predictive ability of five GP models using the genomic best linear unbiased predictor model (GBLUP) with additive + dominance effects (M1) as the baseline and then adding genotype by environment interaction (G × E) (M2), enviromic data (W) (M3), W+G × E (M4), and finally W+G × W (M5), where G × W denotes the genotype by enviromic interaction. Moreover, we considered single-trait multi-environment trials (STMET) and MTMET for three traits: grain yield (GY), plant height (PH), and ear height (EH), with two datasets and two cross-validation schemes. Afterward, we built two kernels for genotype by environment by trait interaction (GET) and genotype by enviromic by trait interaction (GWT) to apply genetic algorithms to select genotype:environment:trait combinations that represent 98% of the variation of the whole dataset and composed the optimized training set (OTS). Using OTS based on enviromic data, it was possible to increase the response to selection per amount invested by 142%. Consequently, our results suggested that genetic algorithms of optimization associated with genomic and enviromic data efficiently design optimized training sets for genomic prediction and improve the genetic gains per dollar invested.

## 1. INTRODUCTION

In the last decades, maize (*Zea mays* L.) has reached the level of the world’s largest crop, being the only one to produce more than 1 billion tons per year (Contini et al., 2019), which makes it a crop of high economic importance, due also to its multiple uses, such as human and animal nutrition, ethanol fuel production and in the pharmaceutical industry. Although maize yield has been growing, the development of new cultivars adapted to the specific edaphoclimatic conditions of different regions at different planting times is still necessary (Andrade et al., 2016).

As an allogamous species of great agronomic interest, maize has already been extensively studied by breeding programs, and the increase in productivity in this species is mainly dependent on the development of single-cross cultivars, also known as hybrids, where the hybridization is used to explore the expression of heterosis, first described by Shull (1908), which is quite expressive and well known in maize. To released new cultivars capable of high yields and great performance of other agronomical characteristics, maize breeding programs develop thousands of hybrids each year that need to be evaluated in field experiments; however, resources are limited, and evaluations are expensive and labor-intensive. The time, area, labor, and budget required to evaluate all those materials in all the desired locations and for all the traits of interest each year are very high and, in most cases, unfeasible. Therefore, technologies, such as genomic prediction (GP), capable of predicting the performance of those materials early, with no need to wait until the end of the crop cycle to discard unwanted materials, are of great interest to the sector (Schrag et al., 2009; Werner et al., 2020).

GP emerged with the promise of increasing genetic gain per unit of time and reducing costs (Meuwissen et al., 2001), and has been widely studied for different crops like maize, wheat, rice, coffee, and brachiaria (Crossa et al., 2017; Carvalho et al., 2020; Matias et al., 2019), as well as for livestock and forest trees. Genomic selection (GS) has been used for many purposes, for example, to predict the performance of lines and double haploids during the initial stages of development (Krchov & Bernardo, 2015; Werner et al., 2020), including the quality (Ibba et al., 2020; Lado et al., 2018), resistance to diseases (Rutkoski et al., 2012) and performance of single-crosses (Bandeira e Sousa et al., 2017; Lyra et al., 2017; Alves et al., 2019). Several groups have shown that there are advantages with the inclusion of multiple traits (MT) in GP, since they explore the correlation between traits and their heritability in the prediction process, surpassing single-trait models’ predictive ability (Jia & Jannink, 2012; Lado et al., 2018; Schulthess et al., 2018). Moreover, the use of multi-environment models (MET) seems unquestionable (Guo et al., 2020; Oakey et al., 2016). Consequently, the combination of both, i.e., multi-trait multi-environment models (MTMET), may improve the accuracy and save labor costs (de Oliveira et al., 2020; Montesinos-López et al., 2016, 2019; Wang et al., 2018).

However, regardless of the GP method, the training set population (TRN) needs to be genotyped and high-quality phenotyped, while the testing set population (TST) only needs to be genotyped. The establishment of the TRN, which should be representative in terms of size, diversity, and the relationship of the individuals to be predicted, is the key to success in GS (Jannink et al., 2010; Akdemir et al., 2015; Crossa et al., 2017; Varshney, 2017; Ibba et al., 2020). For that, the main objectives are to minimize costs associated with phenotyping by selecting smaller training populations, and maximize the predictive ability for the individuals of the TST through efficient resource allocation (Isidro et al., 2015; Lado et al., 2018; Pinho Morais et al., 2020; Riedelsheimer & Melchinger, 2013; Technow et al., 2014). Additionally, there is a lack of knowledge on how to distribute genotypes optimally in multi-environment trials in order to achieve the best balance between the number of genotypes tested in the field and the predictive capacity of GP models, and maximize the selection gain with fixed area and budget resources (Jarquin et al., 2020). Furthermore, in MTMET, we could also ask: which traits should be evaluated in each genotype:trial combination?

Hence, the strategy is to design optimized populations for GP, which allows keeping the accuracy of prediction at satisfactory levels using a training population that is smaller, but representative in terms of information (Fritsche-Neto et al., 2018). In this context, many studies have been carried out aiming to establishing the balance between investment and efficiency through different methods, experiment design, statistical analysis, and TRN composition. For instance, the genetic algorithm to design training populations developed by Akdemir (2017) was tested by Pinho Morais et al. (2020) for several population sizes. The responses were compared with randomly selected populations, noting that optimizing TRN can be effective to obtain satisfactory accuracies. Using MT models, Lado et al. (2018) tested other resource allocation strategies by comparing the PA with different levels of availability of phenotypic information for the target trait (expensive and labor-intensive). For that, they decreased the TRN sizes from 80 to 10%, then included the phenotypic data of all individuals for correlated traits (less laborious and less costly), and finally, considered balanced and unbalanced scenarios. The results showed no loss in PA when reducing TRN for a target trait up to 30% but using full information of correlated traits; additionally, the unbalanced phenotyping approach for correlated traits performed better than the balanced one for the same purpose of reducing TRN. Another strategy was proposed by Costa-Neto et al. (2021a), who investigated the inclusion of dominance effects and envirotyping data into a single-trait MET scenario. The authors found that, especially for traits with low heritability and highly influenced by the environment, the environmental covariables (EC) can increase PA for new environments or newly developed hybrids by tracking variation sources, environment resources, and reducing the error variance.

As described above, the use of accurate genetic algorithms for optimizing training populations can help to reduce the number of genotypes that compose the TRN, as well as reduce costs and field labor, while maintaining good values of predictive accuracy (Akdemir, 2017; Akdemir et al., 2015; Misztal et al., 2014; Misztal, 2016). Additionally, the collection and processing of environmental data can help in the optimization process. Instead of using a simple incidence matrix of environments to model the G × E interaction, processed environmental data better describe specific relationships between environments and crop phenology, called envirotyping. Through envirotyping, it is possible to describe the quality of an environment and estimate the resources available to satisfy the crop needs. When it comes to multi-environment trials (MET), environmental quality ends up as a global average of the entire experimental network. With the aid of some tools, such as the EnvRtype R package of Costa-Neto et al. (2021b), it is possible to compose a covariance matrix (W) between trials, which then makes it possible, among other things, to dissect the G × E interaction, and to build environmental relationship matrices for genomic prediction, which better explain the sources of non-genetic variation, such as the influence of environments on phenotypic variation. Finally, the envirotyping information can be associated with genomic data in genetic algorithms to better select genotypes and target environments that are more informative in terms of G × E (Costa-Neto et al., 2021a).

Compiling these ideas, to optimize TRN sets, MT models may help predict quantitative target traits based on correlated characteristics. MET models also allow the inclusion of the G × E interaction term, which undoubtedly helps predict non-phenotyped individuals. Finally, envirotyping is an emerging component for selecting fewer but well-optimized trial locations. Therefore, our goal was to test the performance of optimized training sets (OTS) for multi-trait multi-environmental trials (MTMET), and the use of environmental covariables (W) in genomic prediction models, with the aim of diminishing the phenotypic labor due to lower but optimally selected population sizes, while keeping the predictive ability at satisfactory levels, and then compare these results with benchmarks. For that, we (i) fitted and compared the performance of five different prediction models, progressively including environmental covariables and interaction terms (G × E and G × W); (ii) estimated the genomic prediction ability of the five prediction models for STMET and MTMET, to use as benchmarks values; and (iii) estimated the genomic prediction ability using OTS with controlled unbalancing of G, E and trait information, selected by a genetic algorithm.

## 2. MATERIALS AND METHODS

### 2.1. Plant material

The phenotypic data consisted of two datasets of tropical maize single-cross hybrids. Plant material was evaluated for the following three traits of agronomic interest: grain yield (GY, in ton ha^-1^), plant height (PH, in cm), and ear height (EH, in cm). For GY assessment, ears were harvested at physiological maturity, grains were adjusted to 13% moisture, and the yield was corrected by area and plant population. PH and EH were measured from the soil surface to the flag leaf collar and the highest ear, respectively, on five representative plants within each plot.

#### HEL dataset

Provided by Helix Seeds (HEL), the first dataset was composed of phenotypic and genotypic data of 452 maize hybrids obtained from single crosses in a partial diallel mating design among 106 tropical maize inbred lines. In order to balance the data, only genotypes that were evaluated in all locations for all traits were considered, so that 247 remained for analysis. Balancing the data will later allow the creation of controlled imbalances. The experimental design used was randomized complete blocks with two replications per genotype per location. Hybrids were evaluated in trials carried out over the 2014/15 growing season at three locations in Brazil: Ipiaçu (IP) and Pato de Minas (PM) in the state of Minas Gerais, and Sertanópolis (SE) in the state of Paraná.

#### USP dataset

The second dataset belongs to the University of Sao Paulo (USP). The data consist of 903 maize single crosses obtained from a diallel mating design between 49 inbred lines. After balancing the data, 623 genotypes remained for analysis. Hybrids were evaluated at two locations in Brazil: Piracicaba (PI) and Anhumas (AN), in São Paulo. They were evaluated for two years during the second growing season of years 2016 and 2017. The experimental design was an augmented block, with two commercial hybrids as checks per block. Although the areas are relatively close on the map, the soil and climate conditions are quite contrasting, and thus characterize different environments, allowing us to consider each location × year combination as an environment: AN.16, PI.16, AN.17, and PI.17.

Further details about both datasets can be found in Alves et al. (2019), Bandeira e Sousa et al. (2017) and Lyra et al. (2017).

### 2.2. Genotypic data

Parental inbred lines from HEL and USP datasets were genotyped with an Affymetrix® Axiom® Maize Genotyping SNP array of 616 K (Unterseer et al., 2014). The genomic quality control (QC) was performed using the SNPRelate package (Zheng et al., 2012) from R software. Markers with a call rate ≤ 0.95 for HEL and a call rate ≤ 0.90 for USP, heterozygous loci in at least one of the parental lines, and monomorphic loci were removed.

The genotypic data of the hybrids were obtained by combining the homozygous markers of their parental lines. The imputation of the lines and genotypes was performed by Synbreed (Wimmer et al., 2012) using the Beagle 4.0 algorithm (Browning & Browning, 2008). Allele frequencies and linkage disequilibrium were computed using the genotypes of the hybrids. Then, markers with minor allele frequency (MAF) ≤ 0.05 were removed. After QC, 30,467 and 62,409 high-quality SNPs were available to analyze the HEL and USP datasets, respectively. All the analyses were performed in the R software (R Core team, 2020).

### 2.3. Enviromic data

Enviromental covariables (EC) were obtained from the EnvRtype R package (Costa-Neto et al., 2021b), to be used as descriptors of the environment for prediction purposes, aiming to increase predictive accuracy (PA) in multi-environment GP scenarios. EnvRtype is a very practical package to acquire and process weather data. Based on trial network information like geographical coordinates (WGS84), plant date, and harvest date, the package collects and processes remote weather data from NASAPower. The environmental factors can be summarized according to the plant phenology intervals of growth or preestablished fixed time intervals. For this research, we used five time intervals according to the maize cycle phenology, defined as 0-14, 15-35, 36-60, 61-90, and 91-120 days after emergency. The environmental factors used were: radiation-related (sunshine hours, in hours, and total daylength, in hours), radiation balance (insolation incident on a horizontal surface, shortwave, and downward thermal infrared radiative flux, longwave), and atmospheric demands (rainfall precipitation, in mm, and relative air humidity, in %) as described in Costa-Neto et al. (2021a). The ECs can be estimated from mean air temperature and accumulated precipitation over the period, for example, and then used to establish G × E interaction. This process creates a covariate matrix of ECs called W, which produces environmental relationship matrices for genomic prediction. Then we can calculate an enviromic kernel equivalent to a genomic relationship matrix, as follows (Costa-Neto et al., 2021b):

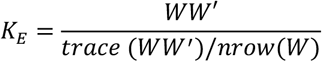

where *K*_*E*_ is the enviromic-based kernel for the similarity between environments and W is the matrix of ECs.

For the HEL dataset, each environment was characterized by 217 ECs, and for the USP dataset, each environment was characterized by 238 ECs, resulting in matrices of dimensions 3 × 217 and 4 × 238, then used to estimate the W matrix.

### 2.4. Variable transformation

We established an index for EH that represents the distance from the actual EH to an ideal ideotype, defined here as 80 centimeters, according to the following formula:

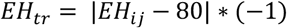

where *EH*_*tr*_ is the transformed EH and *EH*_*ij*_ is the EH for genotype *i* at environment *j*. According to this index, the closer to zero, the closer to our ideal height. For PH, values were normalized in order to obtain a normal distribution interval. To fit the models, all phenotypic data were centered and standardized.

### 2.5. Statistical analysis

#### 2.5.1. Phenotypic analysis

We used a linear mixed model for the two-step analysis to calculate the best linear unbiased estimates (BLUEs) of each trait’s hybrids. BLUEs were obtained within environments for the USP and HEL datasets by the following respective models:

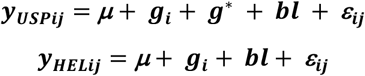

where ***y***_***ij***_ is the estimated phenotypic value of genotype *i* at environment *j*, ***μ*** is the general mean or intercept, ***g***_***i***_ is the fixed effect of hybrid genotype *i*, ***g***^***\****^ is the fixed effect of check genotypes, ***bl*** is the random effect of blocks for the USP dataset 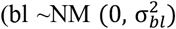 and the fixed effect of blocks for the HEL dataset, and finally, ***ε***_*ij*_ is the residual error for genotype *i* at environment *j*, where ε ∼*N*M(0,*σ*^2^).

Phenotypic analyses were performed using the ASReml-R package (Butler, 2018) of R software (R Core Team, 2020) and subsequently used in our genomic prediction models.

The variance components estimated for each model’s effect will be used to estimate the average broad-sense heritability H^2^.

#### 2.5.2. Genomic prediction scenarios

In order to obtain a benchmark value of PA for the models, we first tested these models in full single-trait multi-environment trials (STMET) and multi-trait multi-environment trials (MTMET) genomic prediction analyses. From those, we were able to obtain the highest possible PA for the specific datasets under study because we used all the information we had available (Fritsche-Neto et al., 2018), through cross-validation schemes with replication.

The algorithms APY (Misztal et al., 2014; Misztal, 2016) and LA-GA-T, from the STPGA R package (Akdemir, 2017), were used in optimization scenarios. The APY is used for determining the size of samples by singular value decomposition. LA-GA-T is a genetic-based algorithm used to select representative individuals from the population and compose the samples. For this purpose, two different kernels were built from the Kronecker product between the variance-covariance matrices of genotypes (G), environments (E), environmental covariables (W), and traits (T) as follows: Σ_G_ ⨂ Σ_E_ ⨂ Σ_T_ and Σ_G_ ⨂ Σ_W_ ⨂ Σ_T_, hereafter called GET and GWT, respectively.

These kernels, used as inputs for the algorithms, assemble combinations between our variables. Thus APY gives us the number of components that explain 98% of the variation within the population, and LA-GA-T selects that number of representative information inside the kernels. Moreover, a genotype was added as a check and therefore evaluated in all environments to create a connection between environments.

The optimized samples from LA-GA-T were obtained three times for each dataset and considered the training set (TRN), while the remaining individuals were used as a testing set (TST). From these three samples (OTS 1), two other scenarios were created, always within kernels. The former one was created by combining the samples two by two (OTS 2), which resulted in three replicates. In the latter, the three independent samples were added together (OTS 3), resulting in just one and bigger optimized training set (OTS).

#### 2.5.3. Genomic prediction via single and multi-trait multi-environment models with additivity and dominance effects

The genomic prediction was first performed by five GBLUP additive + dominance models for STMET and MTMET scenarios. The following models were already tested (for further details, see Costa-Neto et al., 2021a).

##### Model 1 (M1): Environment and main additive plus dominance genomic effects (EAD)

M1 is the most basic model tested, described as follows:

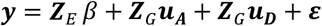

where ***y*** is the adjusted observed values (BLUEs) obtained from the first step for the hybrids. The fixed effects of environment were modeled by ***Z***_***E***_ *β* with the incidence matrix ***Z***_***E***_, and ***Z***_***G***_ is the incidence matrix for the genotypic effects. ***u***_***A***_ is the vector of additive genetic effects, where 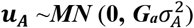, ***u***_***D***_ is the vector of dominance effects, where 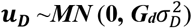, and *ε* is the random residual effect, where 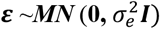. ***G***_***a***_ and ***G***_***d***_ are the genomic relationship matrices (GRM) for additive and dominant effects, respectively, given according to VanRaden (2008) as follows:

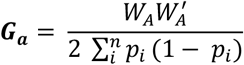

where the values from the incidence matrix ***W***_***A***_ are equal to *0, 1* and *2*, for genotypes markers of *A*_*1*_*A*_*1*_, *A*_*1*_*A*_*2*_ and *A*_*2*_*A*_*2*_, respectively, and *p*_*i*_ is the frequency of one allele from *i* locus.

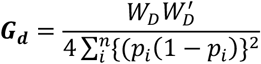

where ***W***_***D***_ contains the values equal to 0 (zero) for both homozygotes *A*_*1*_*A*_*1*_ and *A*_*2*_*A*_*2*,_ and equal to 1 for heterozygotes *A*_*1*_*A*_*2*_.

##### Model 2 (M2): Environment, main effects plus block diagonal GE (EAD+GE)

This model is an update of M1 that accounts for the main effects (A and D), adding the additive × environment and dominance × environment interactions effects (AE and DE).

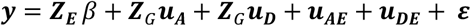

where ***u***_***AE***_ and ***u***_***DE***_ are the vectors of random effects of the interactions. ***u***_***AE***_ and ***u***_***DE***_ have a multivariate normal distribution, 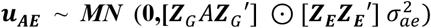 and 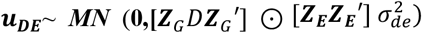, where 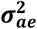 and 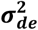 are the variance components for ***u***_***AE***_ and ***u***_***DE***_ interaction effects, respectively (Bandeira e Sousa et al., 2017; Jarquín et al., 2014; Lopez-Cruz et al., 2015).

#### Model 3 (M3): Main effects plus main environmental covariable information (EAD+W)

This third model includes environmental covariables information (W) from envirotyping data.

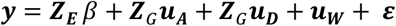

where ***u***_***W***_ is the matrix of environmental covariables, as according to Costa-Neto et al. (2021a), it is non-genetic information that fills the gap between the genomic phenotypic information that remains across environments.

##### Model 4 (M4): Main effects EADW plus reaction norm for GE (EAD+W+GE)

This model is an extension of the previous model (M3), adding the environment’s additive and dominance interactions.

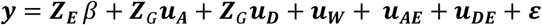

##### Model 5 (M5): Main effects EAD plus W plus reaction norm for GW (EAD+W+GW)

This model is a modification of the latter (M4) reaction-norm variation; it replaces the genomic × environment interactions with the genomic × enviromic effects interactions.

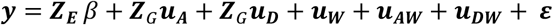

where ***u***_***AW***_ and ***u***_***DW***_ are the vectors of random effects of interactions. ***u***_***AW***_ and ***u***_***DW***_ have a multivariate normal distribution, 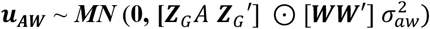 and 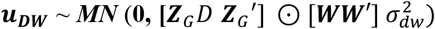. Here we can assume that there are different levels of relationship between genotypes and environments.

All models were fitted with the Bayesian Generalized Linear Regression BGLR R package (Pérez & de los Campos, 2014a; Pérez & De Los Campos, 2014b), using a Gibbs sampler with 10,000 iterations, assuming a burn-in of 1,000, and a thinning of 2.

It is important to point out that the “Multitrait” function of the BGLR package has some basic premises that must be met for the model to work, one of which is the availability of complete information from at least one genotype. Here, as we used multiple traits with the multiple environments approach, we established a genotype as a check, with complete phenotypic data available, common to all environments. This way, it was possible to connect the environments, especially when we explored the G × E interaction.

### 2.6. Assessing the predictive ability of the models

Two cross-validation schemes were used to access GP models’ predictive ability, proposed by Burgueño et al. (2012).

The first validation scheme, known as CV1, was applied considering 50 random partitions with 70% of phenotypic and genotypic (genotypes phenotyped for all traits in all environments) information as TRN, while the remaining 30% (genotypes not phenotyped in any of the environments) were predicted, using only their genotypic information. This scheme aims to quantify GP models’ ability and reproduce a scenario frequently faced by breeders when predicting new genotypes in a network of already known environments, i.e., newly developed maize hybrids never evaluated in any environment. The second scheme, CV2, mimics another common situation when genotypes are tested in unbalanced field trials (or incomplete field trials), i.e., some genotypes are evaluated in some environments but not in the entire experimental network. For this scheme, we also used 50 random partitions with 70% of the information (genotype–environment combinations) as TRN, and the remaining 30% as TST.

For each TRN-TST partition, models were fitted using the TRN, and we performed Pearson’s correlation coefficient between the predicted value and the observed value or BLUE of the TST individuals within each environment, for each one of the 50 partitions. Then these correlations were used to assess the accuracy and compare the performance of each model. Since the BLUEs were calculated by environment, the PAs were also calculated by environment. The same 50 TRN-TST partitions were used to fit each model, allowing access to the best performance model.

For OTS, the predictive ability was also calculated as the Pearson’s correlation coefficient between the predicted value and the adjusted observed value or BLUE of the TST individuals within each environment for each trait; then the average of environments was taken.

### 2.7. Response to selection per unit invested

The genetic gain per dollar invested was estimated to compare the efficiency of the scenarios tested in this work with pure phenotypic selection (PS). The methodology was based on information from Krchov & Bernardo (2015) and Muleta et al. (2019). The phenotyping costs assumed were: 2 US dollars (USD) per plot per trait for PH and EH; 4 USD per plot for GY. For genotyping, we considered 20 USD per sample. As we are dealing with F1 maize hybrids, the parental inbred lines were genotyped, and the hybrid genotype was assembled in silico. This way, the total cost was the sum of the expenses with genotyping (20 USD × number of lines) plus phenotyping the TRN. This calculation was made for each dataset × scenario, considering the three OTS scenarios (OTS 1, OTS 2 and OTS 3) for each kernel and the MTMET CV2 standard scenario. For the phenotypic selection scenario, the average accuracy (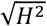 of each trait) was divided by phenotyping cost, 8 USD per plot for all traits, for the complete dataset. The genetic gain was estimated by dividing the PA by the corresponding cost and subsequently transformed to the base of 10,000 USD, given the fact that the other components of the breeder’s equation (Lush, 1937) were considered as fixed.

## 3. RESULTS

### 3.1 Descriptive statistics

Pearson’s correlation between traits was calculated for each dataset using the BLUEs obtained in the first step. As a consequence of the EH transformation, GY assumed a negative correlation with the other traits. For the HEL dataset, GY had a moderately negative correlations with PH and EH, of −0.55 and −0.58, respectively, while PH and EH had a high positive correlation of 0.82. For the USP dataset, GY had weak negative correlations with PH and EH, of −0.44 and −0.33, respectively, while PH and EH had a high positive correlation of 0.70.

Estimated heritability was intermediate to high: for the HEL dataset, trait heritability was 0.62, 0.78 and 0.80 for GY, PH and EH respectively; for the USP dataset, heritability was 0.56 for GY, 0.84 for PH and 0.89 for EH.

As expected, the correlation for the complex trait GY was lower than for PH and EH; additionally, the complex trait had a lower heritability than the auxiliary ones.

### 3.2. Optimized training sets (OTS)

The first result of selecting information to form the training populations, using the APY algorithm, returned the effective population sizes (N_e_) for each kernel, as described below. *HEL dataset:* GET – OTS 1: 155 combined information of genotype × environment × trait selected to form the TRN, which represents 7.4% of observations; GWT– OTS 1: 102 combined information of genotype × environment × trait selected to form the TRN, based on environmental covariables (W), representing 5% of observations. *USP dataset*: GET– OTS 1: 267 combined information of genotype × environment × trait selected to form the TRN set, representing 3.7% of observations; GWT– OTS 1: 107 combined information of genotype × environment × trait selected to form the TRN set, representing 1.6% of observations. Sample size differs depending on the kernel and germplasm because the amount of available information varies as well as the genomic source. From this number, in order to minimize the stochastic error, the LA-GA-T algorithm was performed three times to select the individuals. Thus, the first validation scheme was done for OTS 1; the second combining the three basic populations, two by two, also resulting in three different repetitions, where for Helix the N_e_ were: GET – OTS 2 = 306, representing 13.8% and GWT – OTS 2 = 206, representing 9.3% of total observations, respectively, and for USP: GET – OTS 2 = 533, representing 7.1% and GWT – OTS 2 = 224, representing 3% of total observations, respectively. Finally we added the three repetitions of the base population to form a larger, but optimized, training population, which corresponds to Helix: GET – OTS 3 = 436, representing 19.6% and GWT – OTS 3 = 300 representing 13.5% of total observations, and for USP: GET – OTS 3 = 775, representing 10.4% and GWT – OTS 3 = 326, representing 4.4% of total observations, respectively. The difference in selecting information between the three repetitions from the different OTS tested scenarios can be seen in the heatmaps for Helix (**Fig. 1**a-f) and USP (**Fig. 2**a-f).

**Fig. 1.**
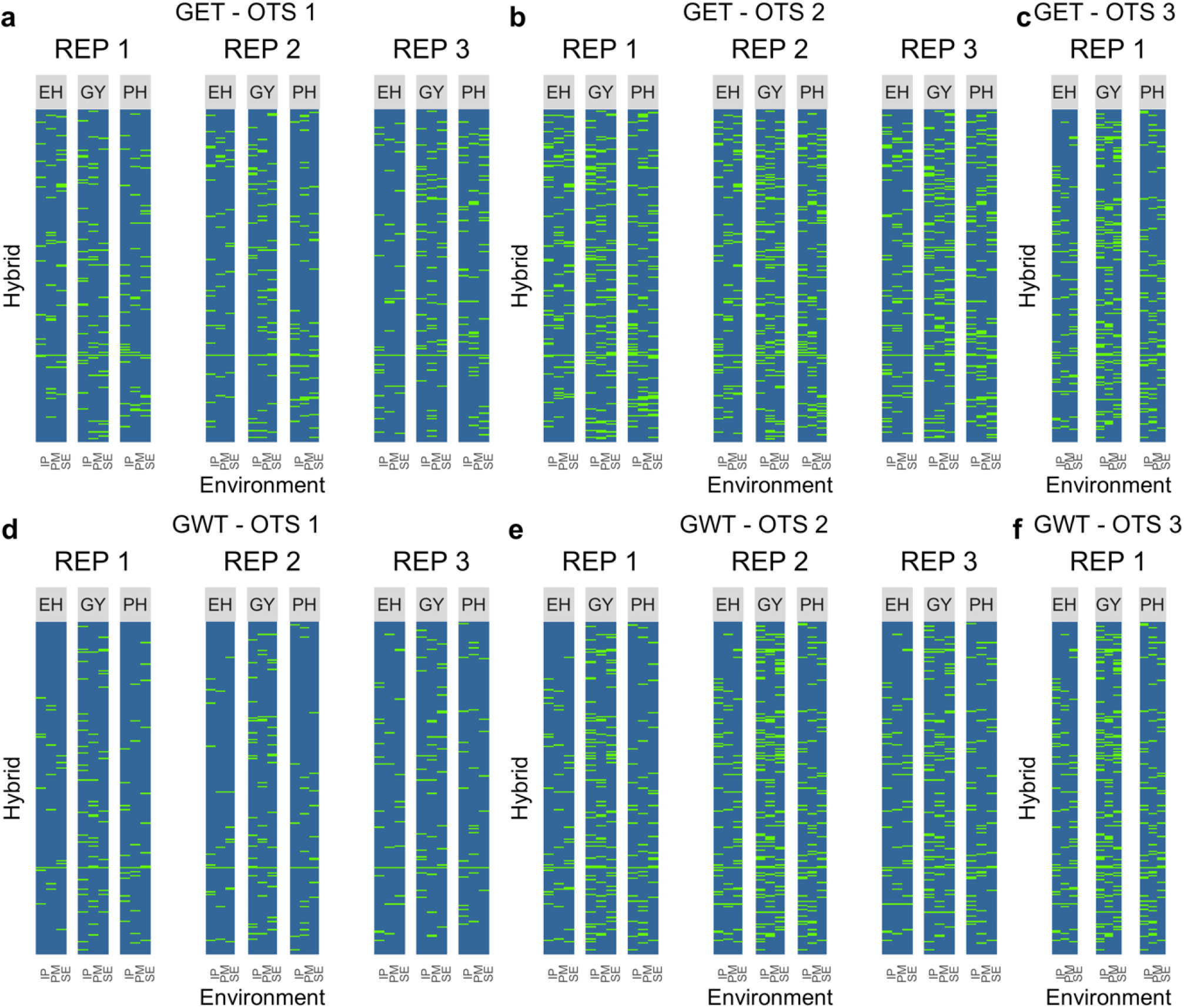
Heatmap of OTS (optimized training sets) graph for the Helix dataset. (a) OTS 1 for kernel GET. (b) OTS 2 for kernel GET. (c) OTS 3 for kernel GET. (d) OTS 1 for kernel GWT. (e) OTS 2 for kernel GWT. (f) OTS 3 for kernel GWT. In green are the hybrids selected to form the training population, for each trait × environment and repetition inside kernels. The solid line that crosses all the graphs represents the genotype used as a check. The environments on the x-axis: IP (Ipiaçu), PM (Patos de Minas), and SE (Sertanópolis); the traits under study: EH (ear height), GY (grain yield), and PH (plant height). The kernels: GET (genotype × environment × trait) and GWT (genotype × environmental covariables × trait) used as the base to select information

**Fig. 2.**
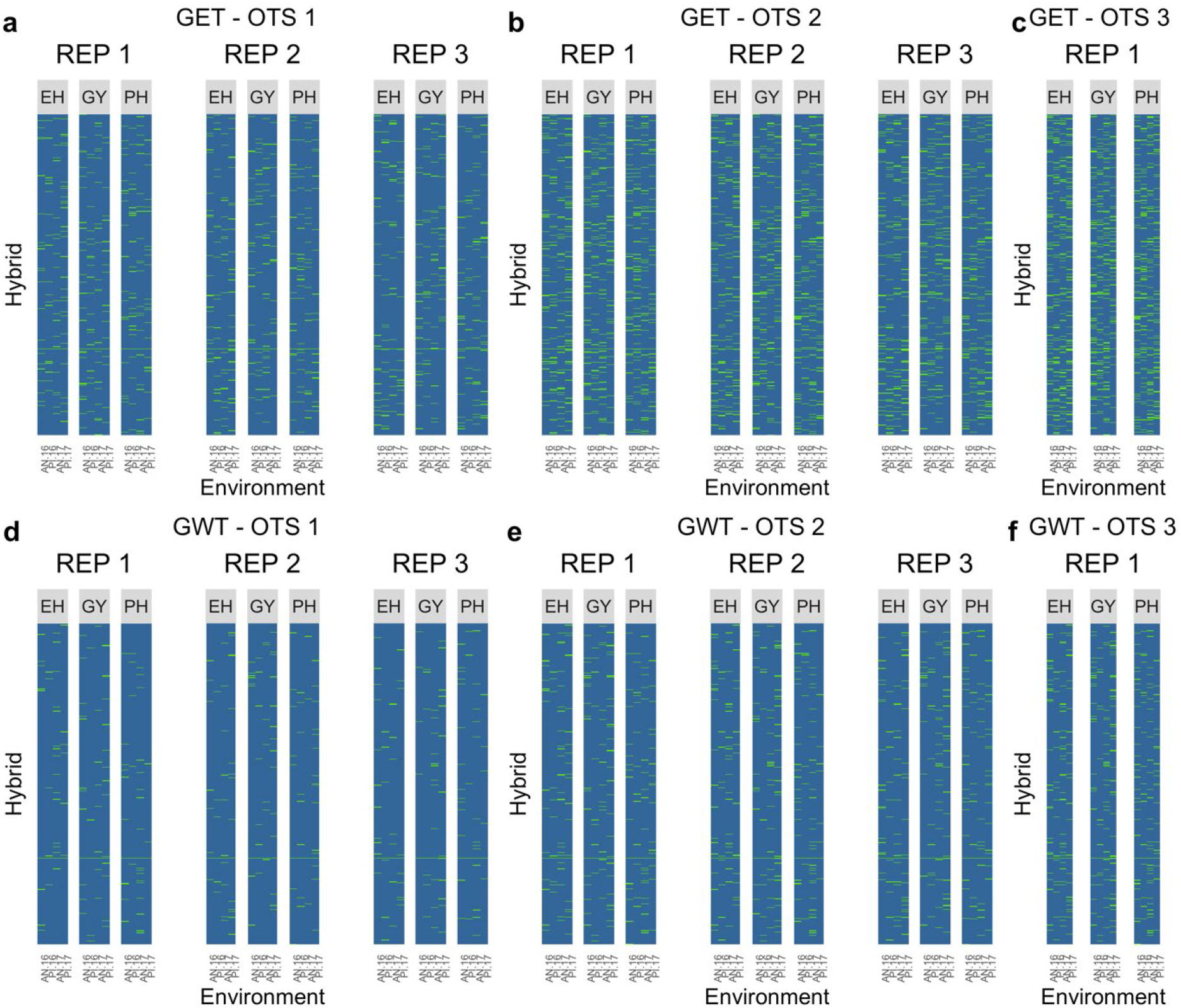
Heatmap of OTS (optimized training sets) graph for the USP dataset. (a) OTS 1 for kernel GET. (b) OTS 2 for kernel GET. (c) OTS 3 for kernel GET. (d) OTS 1 for kernel GWT. (e) OTS 2 for kernel GWT. (f) OTS 3 for kernel GWT. In green we see the distribution of the hybrids selected to form the training population, for each trait x environment and repetition inside kernels. The solid line that crosses all the graphs represents the genotype used as a check. The environments on the x-axis: AN.16 (Anhembi 2016), PI.16 (Piracicaba 2016), AN.17 (Anhembi 2017) and PI.17 (Piracicaba, 2017); the traits under study: EH (ear height), GY (grain yield) and PH (plant height). The kernels: GET (genotype × environment × trait) and GWT (genotype × environmental covariables × trait) are used as the base to select information

### 3.3. Predictive abilities of the five models over STMET and MTMET scenarios HEL dataset

#### Single-trait multi-environment trial analysis

Results for CV1 showed, on average, PAs varying from 0.53 to 0.73 (**Supplementary Table 1**). However, individually, environment SE was inferior in predicting GY, mainly when models without G × E interactions (**M1** and **M3**) were used. In contrast, models including G × E (**M2** and **M4**) increased PA from 100 to 104% and G × W (M5) increased PA by 92% for this specific trait × environment; considering the average within environments, PA for GY between models varies from 3 to 19%. For EH and PH, PAs were higher, from 0.70 to 0.73, and differences between models were minimal, from 0 to 2%. Results for CV2 (Supplementary Table 1) produced the same patterns as those for CV1, ranging from 0.52 to 0.79 when the overall average of environments was considered. For GY in the SE environment, PA could increase between 61 to 68% when models with G × W (**M5**) and G × E interactions (**M2** and **M4**) were used, respectively. For EH and PH, the accuracies were similar within models, with a maximum difference of 2%. Comparing CV1 with CV2, PA means increased from 3 to 8%, and differences were higher for PH and EH. In general, **M4** showed the best performance. Overall, the greatest difference was observed for GY, between models **M1** and **M3** (without G × E interaction) and **M2** and **M4 (**with G × E interaction), where **M4** (**M2**) outperformed **M3** (**M1**) by between 25 and 30%, however, M5 also outperformed **M3** (**M1**) by 29%.

#### Multi-trait multi-environment trial analysis

As for the STMET analysis, the results of the multi-trait analysis showed a similar pattern of responses, both for CV1 and CV2, varying from 0.54 to 0.73 and 0.53 to 0.79, respectively (**Supplementary Table 2**). Nevertheless, we noticed that PAs increased between 61 and 68% for CV1 in the SE environment, which showed the lowest PA for GY in MTMET, like for STMET, when not exploring G × W and G × E interaction effects. Considering the average within environments, the PA variation for GY between models was practically identical to that of STME (0-24%). For EH and PH, accuracy varied from 0.70 to 0.73 and 0.75 to 0.79 for CV1 and CV2, respectively, but in general, the means ranged from 0 to 1%, where **M4** performed better. Comparing CV1 with CV2, the PAs increased from 2.4 to 7.7%, with higher PA differences for GY. Comparing STMET with MTMET, PAs rose from 0 to 1.4%, where the differences were higher for GY and nonexistent for PH.

### USP dataset

#### Single-trait multi-environment trial analysis

Results for CV1 showed PAs varying from 0.46 to 0.65. The PI.17 environment isolated was inferior for predicting EH when models without G × E interactions (**M1** and **M3**) were used (**Supplementary Table 3**). In contrast, models with G × E (**M2** and **M4**) could increase the accuracy up to 440% in the PI.17 environment for EH (considering the overall average within environments, PA for EH varied from 0 to 7%) and model with G × W increased PA by 420% for the same environment × trait; other environments performed very well for EH, with PA from 0.66 to 0.80. For GY, PA ranged from 0.42 to 0.51. For PH, PA were high, from 0.63 to 0.68 between environments, with no difference between models. Results for CV2 produced almost the same patterns as those for CV1. For EH in the PI.17 environment, PA could increase by 340% when models with G × E interactions were used and 320% when using G × W. Despite the environment, PI.17 for EH, the accuracy of all other traits and environments was similar within models, with differences of around 0 to 5%. In general, **M4** showed the best performance. Comparing CV1 with CV2, the PA increased from 4 to 8% for GY and PH, respectively.

#### Multi-trait multi-environment trial analysis

As for the STMET analysis, the results of the multi-trait analysis showed similar response patterns, both for CV1 and CV2 (**Supplementary Table 4)**. Nevertheless, PAs increased between 4 and 8%. Also for STMET, the PI.17 environment showed low PA for EH in MTMET, but by exploring G × E, accuracy increased up to 333%. For GY, accuracies varied from 0.42 to 0.52, and for PH, from 0.63 to 0.72, but for both, in general, the means did not vary. Comparing STMET with MTMET, PAs changed from 0 to 0.4%. Including interaction effects (no matter if G × E or G × W) always increased PA.

### 3.4. Predictive ability for OTS scenarios

Similar to what was done previously, for the optimized training sets, the five models were also tested. The model that achieved the best performance was the **M4**, so only this result will be presented. *Helix dataset:* In the overall average of traits, for OTS 1, PAs were 0.55 and 0.41 for kernels GET and GWT, respectively. Those values increased to 0.63 (+15.9%) and 0.50 (+22.6%), then 0.68 (+7.7%) and 0.61 (+21.4%), for OTS 2 then OTS 3, while the maximum PA obtained by the benchmark CV2 was 0.74 (**Table 1** and **Fig. 3**). *USP dataset:* in the overall average of traits, for OTS 1, PAs were 0.41 and 0.24 for kernels GET and GWT, respectively. Following the same pattern, when we increased the size of TRN, those values increased to 0.47 (+15.6%) and 0.30 (+24.4%) for OTS 2, then 0.52 (+9%) and 0.44 (+46.5%) for OTS 3, while the maximum PA obtained by the benchmark CV2 was 0.61 (**Table 1** and **Fig. 4**).

**Table 1.** Average increase, per trait, in predictive ability, according to an increase in the training set (denoted here as OTS 1, 2 and 3), for both GET and GWT kernels, in the Helix and USP datasets. Ne: number of information used as the training set; prediction accuracies for EH: ear height, GY: grain yield and PH: plant height. Values in parentheses show the percentage increase between that value and the value immediately preceding it

|  | OTS | GET |  |  |  | GWT |  |  |  |
| --- | --- | --- | --- | --- | --- | --- | --- | --- | --- |
|  |  | Ne | EH | GY | PH | Ne | EH | GY | PH |
| Helix | 1 | 155 | 0.61 | 0.47 | 0.56 | 102 | 0.43 | 0.42 | 0.38 |
|  | 2 | 306<br>(+97%) | 0.67<br>(+11,2%) | 0.56<br>(+18%) | 0.66<br>(+19,2%) | 206<br>(+100%) | 0.54<br>(+23,5%) | 0.50<br>(+19,8%) | 0.48<br>(+24,3%) |
|  | 3 | 436<br>(+42%) | 0.72<br>(+7%) | 0.60<br>(+7,7%) | 0.72<br>(+8,5%) | 300<br>(+45%) | 0.64<br>(+18,4%) | 0.60<br>(+19,3%) | 0.63<br>(+26,7%) |
|  | OTS | GET |  |  |  | GWT |  |  |  |
|  |  | Ne | EH | GY | PH | Ne | EH | GY | PH |
| USP | 1 | 267 | 0.47 | 0.28 | 0.48 | 107 | 0.22 | 0.19 | 0.32 |
|  | 2 | 533<br>(+91.9%) | 0.52<br>(+12.6%) | 0.34<br>(+19.8%) | 0.56<br>(+16%) | 224<br>(+87.5%) | 0.31<br>(+42.2%) | 0.23<br>(+23%) | 0.36<br>(+12.8%) |
|  | 3 | 775<br>(+45%) | 0.56<br>(+5.9%) | 0.40<br>(+17%) | 0.61<br>(+8.7%) | 326<br>(+43%) | 0.43<br>(+39%) | 0.35<br>(+51.5%) | 0.54<br>(+49.2%) |

**Fig. 3.**
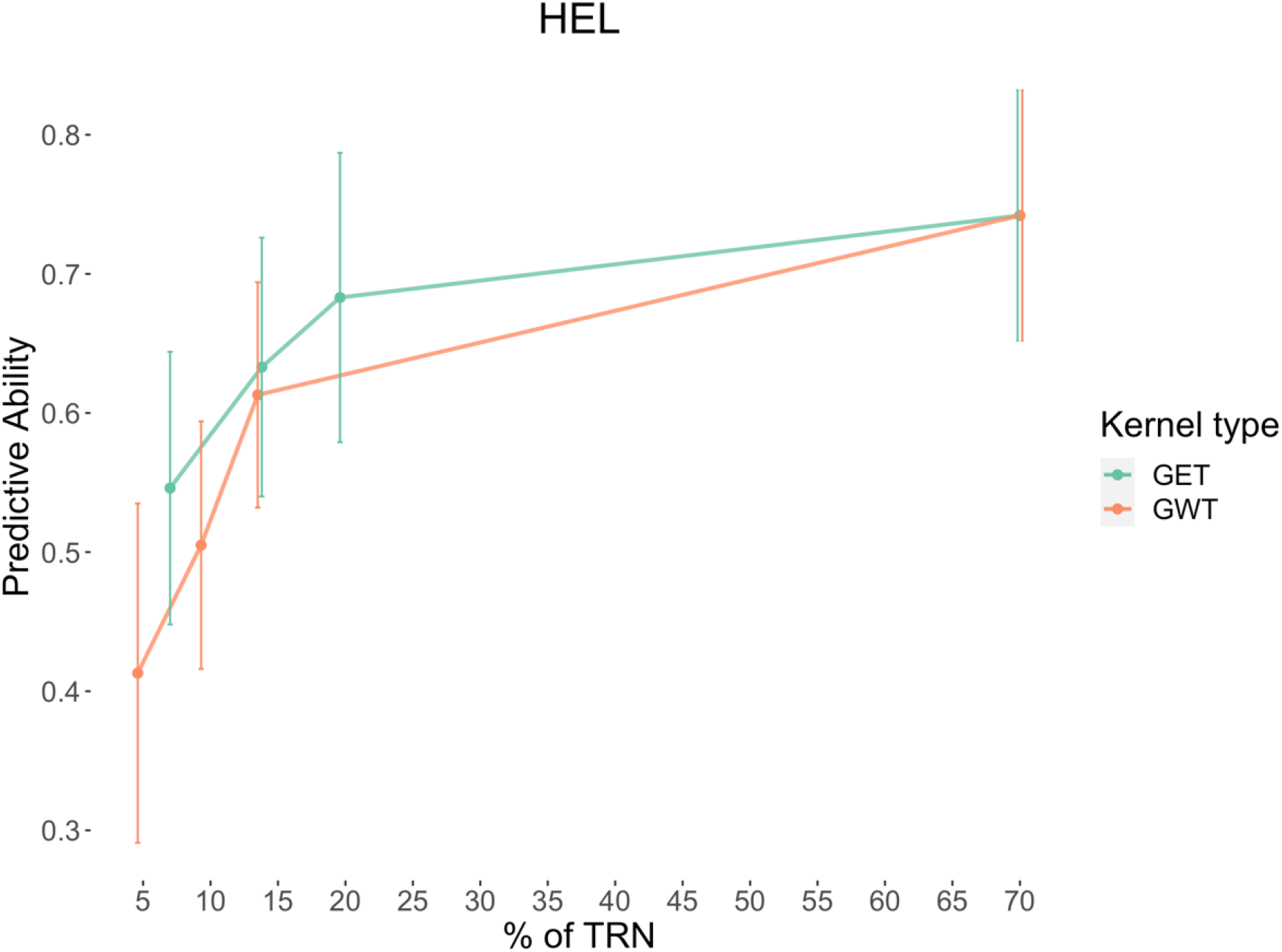
The trend graph shows the increase in the average PA (predictive ability, mean of environments and traits) according to increases in the Helix dataset’s training set size. GET and GWT are the kernels used as the basis for selection of information by the LA-GA-T algorithm. The 4 points on each line correspond to OTS1, OTS2, OTS3 and finally CV2, which is a benchmark or the highest value achieved with this dataset under this particular study’s conditions

**Fig. 4.**
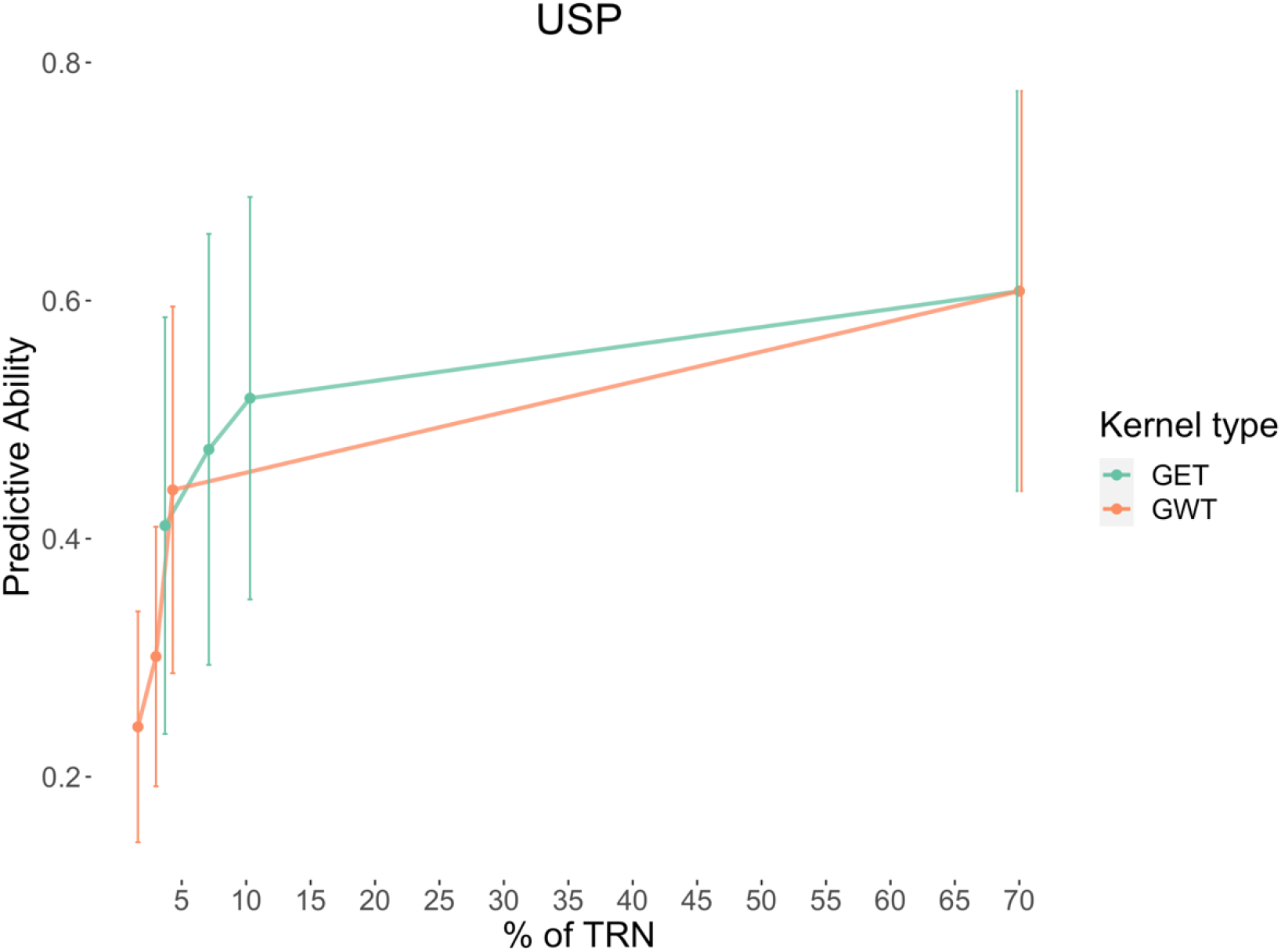
The trend graph shows the increase in the average (predictive ability, mean of environments and traits) according to increases in the USP dataset’s training set size. GET and GWT are the kernels used as the basis for selection of information by the LA-GA-T algorithm. The 4 points on each line correspond to OTS1, OTS2, OTS3 and finally CV2, which is our benchmark or the highest value achieved with this dataset under this particular study’s conditions

### 3.5. Response to selection per amount invested

The genetic gain per dollar per 10,000 USD invested was estimated for each dataset × OTS’ scenario, primarily to compare the efficiency at the kernel level. Then, between scenarios: OTS *versus* MTMET CV2 and OTS *versus* phenotypic selection, within datasets.

For the *HEL dataset*, the results showed an inversion of gain between the kernels over scenarios. While the GET kernel started at 0.80 × 10^−3^ and went to 0.58 × 10^−3^, the GWT kernel started at 0.70 × 10^−3^ and went to 0.67 × 10^−3^ gain per 10,000 USD invested. For PS, the gain was 0.14 × 10^−3^ (**Fig. 5**).

**Fig. 5.**
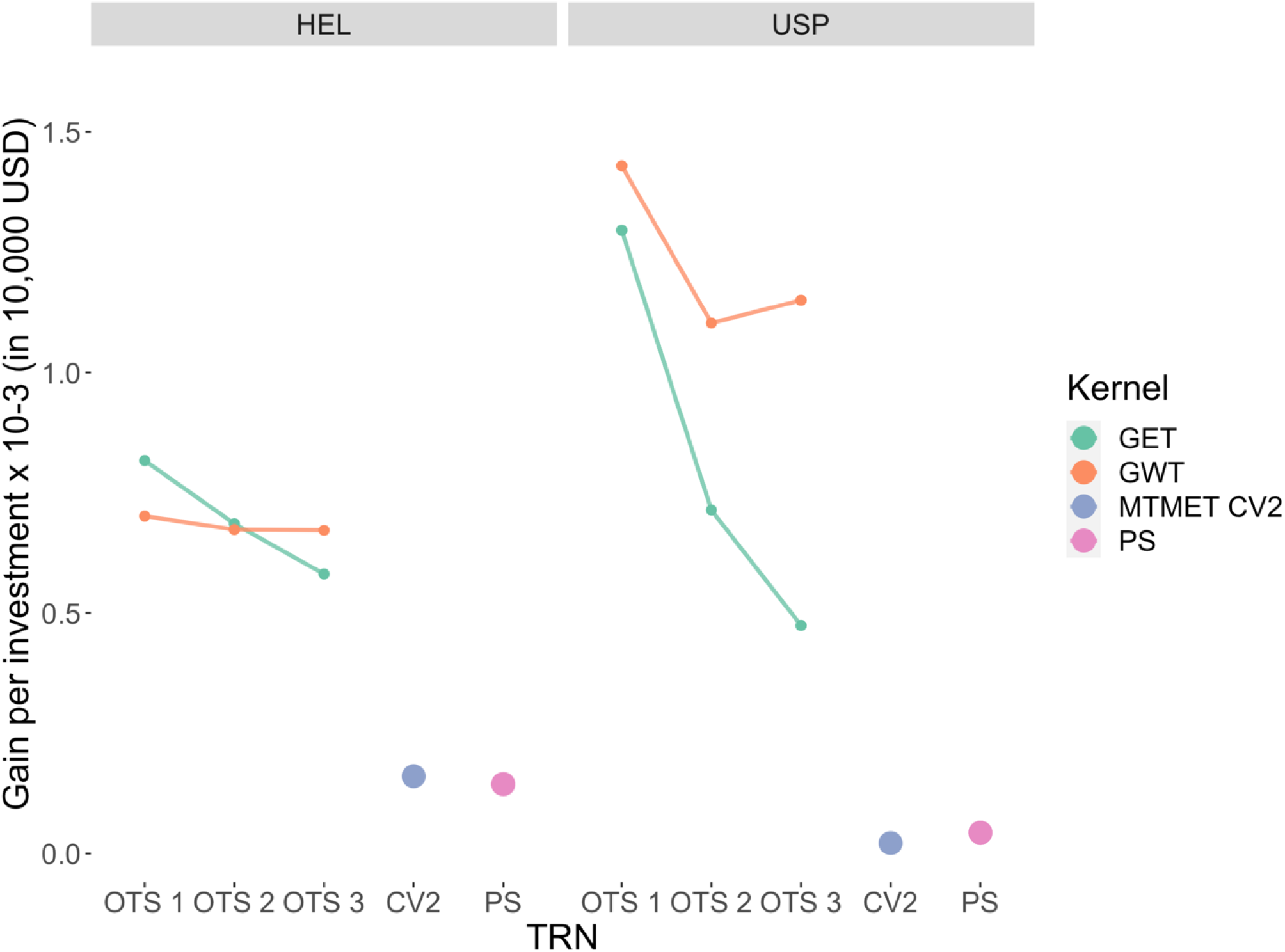
Gain per cost × 10^−3^ (per 10,000 dollars invested) for the HEL and USP datasets, comparing the two optimization kernels (GET and GWT) and the standard scenario (MTMET CV2, TRN = 70%). The cost includes the phenotyping of TRN (3 USD per trait per plot) and the whole dataset’s genotyping (20 USD per sample)

For the *USP dataset*, the GET kernel started with a gain of 1.29 × 10^−3^, and went to 0.47 × 10^−3^, while the GWT kernel started at 1.42 × 10^−3^ and went to 1.15 × 10^−3^. For PS, the gain was 0.04 × 10^−3^.

As a matter of comparison, we did the same procedure for the standard scenario MTMET CV2 (for **M4**) and the gains per investment were 0.16 × 10^−3^ for HEL and 0.02 × 10^−3^ for the USP dataset.

## 4. DISCUSSION

The major goal of GS can be defined as increasing the genetic gain with no increase in costs compared to phenotypic selection only (Crossa et al., 2017; Werner et al., 2020), thus compensating the loss in predictive ability by the gains in response to selection.

The traits evaluated here are moderately to strongly positively correlated by Pearson’s correlation coefficient. However, the selection targets for them in a real breeding program are in opposite directions. While we want to increase grain yield, we want to decrease plant height and stabilize ear height. Dwarf plants with high yield are already a reality in other crops like wheat and rice. However, in maize, dwarfing mutant genes have been studied. Unlike wheat and rice, PH in maize is a quantitative trait that affects other plant characteristics like yield losses, making it difficult to apply (Chen et al., 2018). Since we want all those attributes simultaneously, we created a selection index for EH, which measures the distance from the ideal ideotype defined here as 80 centimeters. Then, traits assume a negative correlation between them due to the index, allowing us to select for all the traits concurrently (Wang et al., 2018).

Similar to what Ibba et al. (2020) and Werner et al. (2020) reported and to what was expected, accuracy is specific to the population, which depends on many other factors like the model chosen, the traits to be predicted, trait heritability, the correlation between traits, the environments, and the correlation between environments. So, the results here were no exception. Differences in predictive ability between the two datasets, which occurred for the benchmarks (**CV1** and **CV2**) and OTS’s kernels (**GET** and **GWT**), can be partially attributed to differences in Pearson’s correlation coefficient between traits, since PA is directly related to the correlation between traits (Lado et al., 2018).

We could see that some models overperformed others, which was also expected since they contain additional terms or variance components such as environmental covariables (**W**) and interaction terms (**G** × **E** and **G** × **W**) that better capture the portion of variance explained by the model (Alves et al., 2019). Results of Dias et al. (2018) suggested that GBLUP models that contain additivity, dominance and G × E interaction should be preferred for predicting the performance of newly developed hybrids in any MET analyses, as is the case with STMET and MTMET under CV1, and OTS.

In this study, two cross-validation schemes, CV1 and CV2 (Burgueño et al., 2012), were used to evaluate PA for both STMET and MTMET models. For all combination scenarios of dataset, model, single or multi-trait, CV2 outperformed CV1, because, in this scheme, we have phenotypic information of genotypes in some environments, but not in others, which helped increase the PA of the models. With CV1 and CV2, single-trait and multi-trait genomic predictions with multi-environment trials are well described and established in the general literature and for the data used in this study (Bandeira e Sousa et al., 2017; Alves et al., 2019). Random cross-validation schemes CV2 and CV1 combined with STMET and MTMET were then used here as benchmarks and as a matter of comparison for the five different prediction models tested (Werner et al., 2020). Thus, based on this prior validation, the model with the highest accuracies (**M4**) was chosen for the prediction using the optimized training set populations, and those validations were also used in a scale as a comparison parameter for prediction accuracies while increasing the effective population sizes of TRN (see **Fig. 3** and **Fig. 4**). Note that the model whose performance was superior includes the G × E interaction component, in agreement with what has already been reported by several authors (Acosta-Pech et al., 2017; Burgueño et al., 2012; Dias et al., 2018; Jarquin et al., 2020; Montesinos-López et al., 2019; Robert et al., 2020) and more recently, by Jarquin et al. (2021). As well, adding the interaction effect G × W (M5) increases PA when compared to the main effects models (M1 and M3), but not as much as models containing the G × E, similar to what was already presented by Robert et al. (2020) and Jarquin et al. (2021); however, G × W has the advantage of predicting new environments (Costa-Neto et al., 2021a; Jarquin et al., 2021; Robert et al., 2020), which has not been tested here. Also, MTMET models gave a small increase in PA compared with STMET, especially for models containing the G × E term (Lyra et al., 2017; Mendonça & Fritsche-Neto, 2020). Furthermore, as Costa-Neto et al. (2021a) reported, including the W matrix helps increase PA, especially in the case of untested hybrids and/or untested environments, by better explaining sources of variation, capturing the environment potential per se and its interaction with the genotypes. Nevertheless, in the present study, and according to what was presented by Jarquin et al. (2021), the inclusion of W matrix alone (M3) did not improve PA, but was similar to the M1; and the inclusion of W with G × E in M4, gave similar results as the G × E alone (M2). Despite that, the W matrix allows optimizing complex information, as we saw in our OTS’s scenarios, then optimizing trials (Jarquin et al., 2021).

Studies involving SNP data, such as GP, require the inverse of the genomic relationship matrix (GRM). However, as the number of individuals to be evaluated increases, the computational cost of this matrix’s inversion is relatively high, with limitations in memory and time. In order to minimize this problem, Misztal (2016) proposed the algorithm for proven and young animals (APY). Using the APY it is not necessary to make a complete inversion of the GRM, since its result returns the number of individuals (n) needed to sample 98% of the population variation; then the inverse of GRM can be obtained by recursion, based on the information of the core individuals. Despite reducing the number of individuals, this algorithm does not specifically indicate which genotypes we should select as core individuals. Hence, it is necessary to take a random sample of population. Here we extended the APY to plants. Since the APY does not provide information on which genotypes should be included in the core population, another algorithm was used to efficiently select these individuals. We used the genetic optimization algorithm in the selection of sub-populations, the LA-GA-T (look ahead genetic algorithm with taboo) proposed by Akdemir (2017) in his STPGA (selection of training populations by genetic algorithms) R package. Genetic algorithms work based on the principles of biological evolution, so that they solve their problems using evolutionary strategies, where at each iteration, the best individuals are selected and elite individuals and so on form the next population. Still, the term taboo indicates that the solutions recently tested will be avoided in the next attempts, avoiding unnecessary evaluations, and limiting the number of iterations necessary to reach convergence. Thus, LA-GA-T optimizes the selection, on a genetic basis, of the *n* genotypes informed by APY to compose our optimized training set (OTS) (Fristche-Neto et al., 2018). In this context, Mendonça & Fritsche-Neto (2020) used the algorithm designed by Akdemir (2017) to select the most representative genotypes to build the training population. Similar to our findings, they did not notice an increase in PA while using OTS, but reduced the budget.

Nevertheless, in the present work, we extended these algorithms to more complex relationship matrices, as the Kronecker product of the genomic relationship matrix (GRM), with the environmental variances and covariances matrix (W) and the traits (T) of interest (G ⊗ E ⊗ T or G ⊗ *W* ⊗ T). Although these scenarios cause a high level of imbalance in the data, they will give an idea about which genotypes need to be phenotyped, in which locations and for which traits in order to form a smaller but optimized training set, reducing fieldwork and the financial resources spent on evaluations, and how much data imbalance the models support for the prediction of hybrids, without important losses in PA. Thus, it allowed us to identify which genotypes should be evaluated in which environments and for which traits, to form a super-optimized training population, capable of predicting the performance of the entire population for all traits and environments, filling gaps in GS and answering questions about the optimal partition of genotypes across environments (Jarquin et al., 2020).

Voss-Fels et al. (2019) said that the TRN have to be exceptionally large. Similarly, Wang et al. (2018) stated that the larger the TRN, the better the estimation of genetic effects and therefore, the greater the accuracy, mainly for low heritability traits. Here, the amount of information for each OTS, regardless of the kernel, is small, representing between 1.6 and 19.6% of the total available information. Moreover, the samples have a good distribution of genotypes, including those genotypes that perform well, and those that are not so good, bringing positive impacts on PA (Michel et al., 2020). As expected and similar to what Pinho Morais et al. (2020) found, with a small effective population size, PA is diminished, since the sample contains small genetic variability. However, we still obtained satisfactory prediction values, with an overall mean up to 0.41 and 0.54 for USP and Helix, respectively, with the advantage that costs were reduced by more than 1000% and the labor with the TRN was also reduced. According to Krchov and Bernardo (2015), accuracies should be greater than 0.50 so that the GS is superior and chosen instead of the phenotypic selection.

It is interesting to notice that, with a similar increase in effective population size of approximately 90-100% from OTS 1 to OTS 2, and from OTS 2 to OTS 3, respecting the particularities of each training set population, the increases in PA are different for each kernel-dataset combination, as given in **Fig. 1**, **Fig. 2** and **Table 1**, which show that PA practically doubled for GWT when compared to GET, mainly for the USP dataset, whose amount of information used as TRN represents a very small portion of the total dataset. Therefore, it means that a small increase in the training population results in satisfactory PA increases, especially when considering the response to selection. However, GET proved less efficient for optimizing TRN, because it assumes that the environments are not related, and thus needs more information to explain variations in the whole dataset, while GWT proved more efficient for optimizing TRN, than using the W matrix for optimization. With all the aggregated information it carries, GWT is able to select individuals more assertively, and then needs less information to form the OTS, so the TRN size is smaller, providing the added advantages of lower cost and labor. Shown here for a complex case, the global idea is similar to what was observed in Costa-Neto et al. (2021a), adding information to help in prediction.

Since the costs of genotyping are decreasing, whereas the costs of field testing in maize are either stagnant or increasing, and adding that genotypic information is stable, not liable to seasonal variation, less effort is expended saving money resources, genotypic selection is, from this perspective, more efficient (Krchov & Bernardo, 2015). It is worth noting that with a fixed budget, as we decrease the training population size, using OTS for example, we can have a larger test population, since the resources are reallocated from phenotyping to genotyping, or even, with a fixed training population, small budget increments mean a significant increase in the test population, which can be considerably expanded for greater selection gains (Riedelsheimer & Melchinger, 2013; Krchov & Bernardo, 2015).

Looking to reduce costs with phenotyping, Lado et al. (2018), working with wheat, found that there were no losses in the predictive power when reducing the training population up to 30% when traits highly correlated to the trait of interest are used in the multi-trait model because the correlation between the traits and the heritability of each assists in the prediction of the other. Here, we could reduce the training population up to approximately 4% obtaining satisfactory results when considering the selection gains per 10,000USD invested, which were about 0.70 × 10^−3^ against 0.16 × 10^−3^, with optimized (**GWT**) against standard scenario (MTMET CV2) TRN, respectively, for the HEL dataset and 1.42 × 10^−3^ against 0.16 × 10^−3^ for the USP dataset.

The genetic gain per dollar invested was estimated as a basis of comparison for responses in different TRN sizes, datasets and mainly to justify the use of optimization for training populations in GS. From the results, we can infer that the optimized populations have advantages over the standard scenario (70% TRN-30% TST). The difference in gains between the datasets is due to their particular characteristics, such as the number of inbred lines and the PA reached. For the HEL dataset, however, PA was higher, and the costs of genotyping were also higher, since there were a great number of inbred lines, so gains were lower; there was also an inversion of gains between GET and GWT, where the efficiency of the GWT kernel remains practically constant while that of GET falls. For the USP dataset it was the opposite; nevertheless, the PAs were lower, there were fewer individuals to both genotype and phenotype. In this way, the costs per individual were lower and the gains were higher, and GWT outperformed GET in all the scenarios. Altogether, GWT allowed reducing the TRN up to 58% compared with GET. From this point of view, although the PA using GWT is the lowest, independent of the scenario, its advantage can be offset by the gain in the response to selection per USD invested (see **Fig. 5**); giving special attention to GWT in the USP dataset. In summary, compared with MTMET – CV2, with GWT there was a reduction of up to 60% in terms of PA; however, it brings the possibility of substantially reducing the number of plot:traits to be phenotyped up to 98%. Furthermore, using OTS plus W allows increasing the response to selection per amount invested up to 142% compared with GET; thus, there is no gain in PA with OTS, but the reduction in the training population greatly reduces costs and fieldwork, and thus, the relative genetic gains are greater.

In light of this, our results add to the subject of training populations, answering questions about which design to use to distribute the population for evaluation, which individuals to choose to form the training population, because as already seen, TRN is the key to the success of GS.

Multi-trait and multi-environment analyses have been applied as a way to optimize the distribution of resources through GP, reaching satisfactory accuracy and gains; however, this scenario can still be worked on and improved, taking advantage of new tools, like the environmental relationship matrix (W matrix) and genetic algorithms (APY and LA-GA-T) to optimize the allocation of resources. To our knowledge, this is the first work that tests the optimization of training set populations with genetic algorithms, which determine the size of the population and select the individuals based on complex kernels, causing a high level of imbalance, and we observed that, using a smaller optimized training set, we diminished the phenotypic evaluations in the field, and consequently saved costs that can be reallocated for genotyping. Additionally, we calculated the gains per 10,000USD invested, which allowed us to infer that, in a practical way, by applying optimization and maintaining a constant selection intensity, under a fixed budget, the input lines/hybrids of a breeding program can be larger, a greater number of crosses can be tested per cycle and in the early stages, this will improve the gains. The initial investments in GS are considerably high; however, they are offset by gains per unit of time. Nevertheless, it is known that that the genetic variance of a given population decreases over the selection cycles, especially in small populations sizes, which limits the selection gains (Muleta et al., 2019). Hence, we can consider the optimization aiming to renovate training sets each year, to keep GS accuracy acceptable and raise the gains. Therefore, periodic recalibration of the training population is important to endorse genetic variability and, in addition, when using an optimized population for recalibration, the cost of evaluation (genotyping plus phenotyping) is offset by the genetic gains obtained (Muleta et al., 2019). In summary, optimization gave a good balance between gain *versus* costs, and between gain *versus* labor, and added new insight for using the algorithms tested here.

## 5. CONCLUSIONS

Genomic prediction models that include G × E and G × W interaction effects always increase PA, performing better than main effects models; G × E interaction is the best scenario, with a small increase in multi-trait multi-environment analysis when compared with single-trait multi-environment analysis. Furthermore, genetic algorithms of optimization associated with genomic and enviromic data are efficient in designing optimized training sets for genomic prediction and improve genetic gains per dollar invested. However, it is worth remembering that there is a specific interaction between combinations of germplasm, environments, experimental network and evaluated traits that must be taken into account when using the proposed approach.

## REFERENCES

Acosta-Pech, R., Crossa, J., de los Campos, G., Teyssèdre, S., Claustres, B., Pérez-Elizalde, S., & Pérez-Rodríguez, P. (2017). Genomic models with genotype × environment interaction for predicting hybrid performance: an application in maize hybrids. Theoretical and Applied Genetics, 130(7), 1431–1440. https://doi.org/10.1007/s00122-017-2898-0

Akdemir, D. (2017). STPGA: Selection of training populations with a genetic algorithm (p. 111989). https://doi.org/10.1101/111989

Akdemir, D., Sanchez, J. I., & Jannink, J. L. (2015). Optimization of genomic selection training populations with a genetic algorithm. Genetics Selection Evolution, 47(1), 1–10. https://doi.org/10.1186/s12711-015-0116-6

Alves, F. C., Granato, Í. S. C., Galli, G., Lyra, D. H., Fritsche-Neto, R., & De Los Campos, G. (2019). Bayesian analysis and prediction of hybrid performance. Plant Methods, 15(1), 1–18. https://doi.org/10.1186/s13007-019-0388-x

Andrade, L. R. B. de, Neto, R. F., Granato, Í. S. C., Sant’Ana, G. C., Morais, P. P. P., & Borém, A. (2016). Genetic vulnerability and the relationship of commercial germplasms of maize in Brazil with the nested association mapping parents. PLoS ONE, 11(10), 1–14. https://doi.org/10.1371/journal.pone.0163739

Bandeira e Sousa, M., Cuevas, J., Couto, E. G. de O., Pérez-Rodríguez, P., Jarquín, D., Fritsche-Neto, R., Burgueño, J., & Crossa, J. (2017). Genomic-enabled prediction in maize using kernel models with genotype × environment interaction. G3: Genes, Genomes, Genetics, 7(6), 1995–2014. https://doi.org/10.1534/g3.117.042341

Browning, B. L., & Browning, S. R. (2008). A unified approach to genotype imputation and haplotype-phase inference for large data sets of trios and unrelated individuals. American Journal of Human Genetics, 84(2), 210–223. https://doi.org/10.1016/j.ajhg.2009.01.005

Burgueño, J., de los Campos, G., Weigel, K., & Crossa, J. (2012). Genomic prediction of breeding values when modeling genotype × environment interaction using pedigree and dense molecular markers. Crop Science, 52(2), 707–719. https://doi.org/10.2135/cropsci2011.06.0299

Butler, A. D. (2018). Package ‘ asreml .’

Carvalho, H. F., Galli, G., Ventorim Ferrão, L. F., Vieira Almeida Nonato, J., Padilha, L., Perez Maluf, M., Ribeiro de Resende, M. F., Guerreiro Filho, O., & Fritsche-Neto, R. (2020). The effect of bienniality on genomic prediction of yield in arabica coffee. Euphytica, 216(6). https://doi.org/10.1007/s10681-020-02641-7

Chen, Q., Song, J., Du, W. P., Xu, L. Y., Jiang, Y., Zhang, J., Xiang, X. L., & Yu, G. R. (2018). Identification and genetic mapping for rht-DM, a dominant dwarfing gene in mutant semi-dwarf maize using QTL-seq approach. Genes and Genomics, 40(10), 1091–1099. https://doi.org/10.1007/s13258-018-0716-y

Contini, E., Mota, M. M., Marra, R., Borghi, E. Miranda, R. A. de Silva, A. F. da Silva, D. D. da, Machado, J. R. de A., Cota, L. V. Costa, R. V. da, & Mendes, S. M. (2019). Milho-Caracterização e Desafios Tecnológicos. Embrapa, 5(1), 1–45.

Costa-Neto, G., Fritsche-Neto, R., & Crossa, J. (2021). Nonlinear kernels, dominance, and envirotyping data increase the accuracy of genome-based prediction in multi-environment trials. Heredity, 126(1), 92–106. https://doi.org/10.1038/s41437-020-00353-1

Costa-Neto, G., Galli, G., Carvalho, H. F., Crossa, J., & Fritsche-Neto, R. (2021). EnvRtype : a software to interplay enviromics and quantitative genomics in agriculture . G3 Genes|Genomes|Genetics. https://doi.org/10.1093/g3journal/jkab040

Crossa, J., Pérez-Rodríguez, P., Cuevas, J., Montesinos-López, O., Jarquín, D., de los Campos, G., Burgueño, J., González-Camacho, J. M., Pérez-Elizalde, S., Beyene, Y., Dreisigacker, S., Singh, R., Zhang, X., Gowda, M., Roorkiwal, M., Rutkoski, J., & Varshney, R. K. (2017). Genomic Selection in Plant Breeding: Methods, Models, and Perspectives. Trends in Plant Science, 22(11), 961–975. https://doi.org/10.1016/j.tplants.2017.08.011

de Oliveira, A. A., Resende, M. F. R., Ferrão, L. F. V., Amadeu, R. R., Guimarães, L. J. M., Guimarães, C. T., Pastina, M. M., & Margarido, G. R. A. (2020). Genomic prediction applied to multiple traits and environments in second season maize hybrids. Heredity, 125(1–2), 60–72. https://doi.org/10.1038/s41437-020-0321-0

Dias, K. O. D. G., Gezan, S. A., Guimarães, C. T., Nazarian, A., Da Costa E Silva, L., Parentoni, S. N., De Oliveira Guimarães, P. E., De Oliveira Anoni, C., Pádua, J. M. V., De Oliveira Pinto, M., Noda, R. W., Ribeiro, C. A. G., De Magalhães, J. V., Garcia, A. A. F., De Souza, J. C., Guimarães, L. J. M., & Pastina, M. M. (2018). Improving accuracies of genomic predictions for drought tolerance in maize by joint modeling of additive and dominance effects in multi-environment trials. Heredity, 121(1), 24–37. https://doi.org/10.1038/s41437-018-0053-6

Fristche-Neto, R., Akdemir, D., & Jannink, J. L. (2018). Accuracy of genomic selection to predict maize single-crosses obtained through different mating designs. Theoretical and Applied Genetics, 131(5), 1153–1162. https://doi.org/10.1007/s00122-018-3068-8

Guo, J., Pradhan, S., Shahi, D., Khan, J., Mcbreen, J., Bai, G., Murphy, J. P., & Babar, M. A. (2020). Increased Prediction Accuracy Using Combined Genomic Information and Physiological Traits in A Soft Wheat Panel Evaluated in Multi-Environments. Scientific Reports, 10(1), 1–12. https://doi.org/10.1038/s41598-020-63919-3

Ibba, M. I., Crossa, J., Montesinos-López, O. A., Montesinos-López, A., Juliana, P., Guzman, C., Delorean, E., Dreisigacker, S., & Poland, J. (2020). Genome-based prediction of multiple wheat quality traits in multiple years. Plant Genome, 13(3). https://doi.org/10.1002/tpg2.20034

Isidro, J., Jannink, J. L., Akdemir, D., Poland, J., Heslot, N., & Sorrells, M. E. (2015). Training set optimization under population structure in genomic selection. TAG. Theoretical and Applied Genetics. Theoretische Und Angewandte Genetik, 128(1), 145–158. https://doi.org/10.1007/s00122-014-2418-4

Jannink, J. L., Lorenz, A. J., & Iwata, H. (2010). Genomic selection in plant breeding: From theory to practice. Briefings in Functional Genomics and Proteomics, 9(2), 166–177. https://doi.org/10.1093/bfgp/elq001

Jarquín, D., Crossa, J., Lacaze, X., Du Cheyron, P., Daucourt, J., Lorgeou, J., Piraux, F., Guerreiro, L., Pérez, P., Calus, M., Burgueño, J., & de los Campos, G. (2014). A reaction norm model for genomic selection using high-dimensional genomic and environmental data. Theoretical and Applied Genetics, 127(3), 595–607. https://doi.org/10.1007/s00122-013-2243-1

Jarquin, D., Howard, R., Crossa, J., Beyene, Y., Gowda, M., Martini, J. W. R., Pazaran, G. C., Burgueño, J., Pacheco, A., Grondona, M., Wimmer, V., & Prasanna, B. M. (2020). Genomic prediction enhanced sparse testing for multi-environment trials. G3: Genes, Genomes, Genetics, 10(8), 2725–2739. https://doi.org/10.1534/g3.120.401349

Jarquin, D. Leon, N. De, Romay, C., Bohn, M., & Buckler, E. S. (2021). Utility of Climatic Information via Combining Ability Models to Improve Genomic Prediction for Yield Within the Genomes to Fields Maize Project. 11(March), 1–11. https://doi.org/10.3389/fgene.2020.592769

Jia, Y., & Jannink, J. L. (2012). Multiple-trait genomic selection methods increase genetic value prediction accuracy. Genetics, 192(4), 1513–1522. https://doi.org/10.1534/genetics.112.144246

Krchov, L. M., & Bernardo, R. (2015). Relative efficiency of genomewide selection for testcross performance of doubled haploid lines in a maize breeding program. Crop Science, 55(5), 2091–2099. https://doi.org/10.2135/cropsci2015.01.0064

Lado, B., Vázquez, D., Quincke, M., Silva, P., Aguilar, I., & Gutiérrez, L. (2018). Resource allocation optimization with multi-trait genomic prediction for bread wheat (Triticum aestivum L.) baking quality. Theoretical and Applied Genetics, 131(12), 2719–2731. https://doi.org/10.1007/s00122-018-3186-3

Lopez-Cruz, M., Crossa, J., Bonnett, D., Dreisigacker, S., Poland, J., Jannink, J. L., Singh, R. P., Autrique, E., & de los Campos, G. (2015). Increased prediction accuracy in wheat breeding trials using a marker × environment interaction genomic selection model. G3: Genes, Genomes, Genetics, 5(4), 569–582. https://doi.org/10.1534/g3.114.016097

Lush, J.L. (1937). Animal Breeding Plans. Collegiate Press, Inc., Ames.

Lyra, D. H., de Freitas Mendonça, L., Galli, G., Alves, F. C., Granato, Í. S. C., & Fritsche-Neto, R. (2017). Multi-trait genomic prediction for nitrogen response indices in tropical maize hybrids. Molecular Breeding, 37(6). https://doi.org/10.1007/s11032-017-0681-1

Matias, F. I., Alves, F. C., Meireles, K. G. X., Barrios, S. C. L., do Valle, C. B., Endelman, J. B., & Fritsche-Neto, R. (2019). On the accuracy of genomic prediction models considering multi-trait and allele dosage in Urochloa spp. interspecific tetraploid hybrids. Molecular Breeding, 39(7). https://doi.org/10.1007/s11032-019-1002-7

Mendonça, L. de F., & Fritsche-Neto, R. (2020). The accuracy of different strategies for building training sets for genomic predictions in segregating soybean populations. Crop Science, 60(6), 3115–3126. https://doi.org/10.1002/csc2.20267

Meuwissen, T. H. E., Hayes, B. J., & Goddard, M. E. (2001). Prediction of total genetic value using genome-wide dense marker maps. Genetics, 157(4), 1819–1829. https://doi.org/10.1093/genetics/157.4.1819

Michel, S., Löschenberger, F., Sparry, E., Ametz, C., & Bürstmayr, H. (2020). Mitigating the impact of selective phenotyping in training populations on the prediction ability by multi-trait pedigree and genomic selection models. Plant Breeding, 139(6), 1067–1075. https://doi.org/10.1111/pbr.12862

Misztal, I., Legarra, A., & Aguilar, I. (2014). Using recursion to compute the inverse of the genomic relationship matrix. Journal of Dairy Science, 97(6), 3943–3952. https://doi.org/10.3168/jds.2013-7752

Misztal, Ignacy. (2016). Inexpensive computation of the inverse of the genomic relationship matrix in populations with small effective population size. Genetics, 202(2), 401–409. https://doi.org/10.1534/genetics.115.182089

Montesinos-López, O. A., Montesinos-López, A., Crossa, J., Toledo, F. H., Pérez-Hernández, O., Eskridge, K. M., & Rutkoski, J. (2016). A genomic bayesian multi-trait and multi-environment model. G3: Genes, Genomes, Genetics, 6(9), 2725–2774. https://doi.org/10.1534/g3.116.032359

Montesinos-López, O. A., Montesinos-López, A., Tuberosa, R., Maccaferri, M., Sciara, G., Ammar, K., & Crossa, J. (2019). Multi-Trait, Multi-Environment Genomic Prediction of Durum Wheat With Genomic Best Linear Unbiased Predictor and Deep Learning Methods. Frontiers in Plant Science, 10(November), 1–12. https://doi.org/10.3389/fpls.2019.01311

Muleta, K. T., Pressoir, G., & Morris, G. P. (2019). Optimizing genomic selection for a sorghum breeding program in Haiti: A simulation study. G3: Genes, Genomes, Genetics, 9(2), 391–401. https://doi.org/10.1534/g3.118.200932

Oakey, H., Cullis, B., Thompson, R., Comadran, J., Halpin, C., & Waugh, R. (2016). Genomic selection in multi-environment crop trials. G3: Genes, Genomes, Genetics, 6(5), 1313–1326. https://doi.org/10.1534/g3.116.027524

Pérez, P., & de los Campos, G. (2014). BGLR : A Statistical Package for Whole Genome Regression and Prediction. Genetics, 198(2), 483–495.

Pérez, P., & De Los Campos, G. (2014). Genome-wide regression and prediction with the BGLR statistical package. Genetics, 198(2), 483–495. https://doi.org/10.1534/genetics.114.164442

Pinho Morais, P. P., Akdemir, D., Braatz de Andrade, L. R., Jannink, J. L., Fritsche-Neto, R., Borém, A., Couto Alves, F., Hottis Lyra, D., & Granato, Í. S. C. (2020). Using public databases for genomic prediction of tropical maize lines. Plant Breeding, 139(4), 697–707. https://doi.org/10.1111/pbr.12827

R Core Team (2020). R: A language and environment for statistical computing. R Foundation for Statistical Computing, Vienna, Austria. URL https://www.R-project.org/.

Riedelsheimer, C., & Melchinger, A. E. (2013). Optimizing the allocation of resources for genomic selection in one breeding cycle. Theoretical and Applied Genetics, 126(11), 2835–2848. https://doi.org/10.1007/s00122-013-2175-9

Robert, P. Gouis, J. Le, Consortium, T. B., & Rincent, R. (2020). Combining Crop Growth Modeling With Trait-Assisted Prediction Improved the Prediction of Genotype by Environment Interactions. 11(June), 1–11. https://doi.org/10.3389/fpls.2020.00827

Rutkoski, J., Benson, J., Jia, Y., Brown-Guedira, G., Jannink, J.-L., & Sorrells, M. (2012). Evaluation of Genomic Prediction Methods for Fusarium Head Blight Resistance in Wheat. The Plant Genome, 5(2), 51–61. https://doi.org/10.3835/plantgenome2012.02.0001

Schrag, T. A., Möhring, J., Maurer, H. P., Dhillon, B. S., Melchinger, A. E., Piepho, H. P., Sørensen, A. P., & Frisch, M. (2009). Molecular marker-based prediction of hybrid performance in maize using unbalanced data from multiple experiments with factorial crosses. Theoretical and Applied Genetics, 118(4), 741–751. https://doi.org/10.1007/s00122-008-0934-9

Shull, G. H. (1908). The Composition of a Field of Maize, Journal of Heredity, Volume os-4, Issue 1, January 1908, Pages 296–301, https://doi.org/10.1093/jhered/os-4.1.296

Schulthess, A. W., Zhao, Y., Longin, C. F. H., & Reif, J. C. (2018). Advantages and limitations of multiple-trait genomic prediction for Fusarium head blight severity in hybrid wheat (Triticum aestivum L.). Theoretical and Applied Genetics, 131(3), 685–701. https://doi.org/10.1007/s00122-017-3029-7

Technow, F., Schrag, T. A., Schipprack, W., Bauer, E., Simianer, H., & Melchinger, A. E. (2014). Genome properties and prospects of genomic prediction of hybrid performance in a breeding program of maize. Genetics, 197(4), 1343–1355. https://doi.org/10.1534/genetics.114.165860

Unterseer, S., Bauer, E., Haberer, G., Seidel, M., Knaak, C., Ouzunova, M., Meitinger, T., Strom, T. M., Fries, R., Pausch, H., Bertani, C., Davassi, A., Mayer, K. F. X., & Schön, C. C. (2014). A powerful tool for genome analysis in maize: Development and evaluation of the high density 600 k SNP genotyping array. BMC Genomics, 15(1), 1–15. https://doi.org/10.1186/1471-2164-15-823

VanRaden, P. M. (2008). Efficient methods to compute genomic predictions. Journal of Dairy Science, 91(11), 4414–4423. https://doi.org/10.3168/jds.2007-0980

Voss-Fels, K. P., Cooper, M., & Hayes, B. J. (2019). Accelerating crop genetic gains with genomic selection. Theoretical and Applied Genetics, 132(3), 669–686. https://doi.org/10.1007/s00122-018-3270-8

Wang, X., Xu, Y., Hu, Z., & Xu, C. (2018). Genomic selection methods for crop improvement: Current status and prospects. Crop Journal, 6(4), 330–340. https://doi.org/10.1016/j.cj.2018.03.001

Werner, C. R., Gaynor, R. C., Gorjanc, G., Hickey, J. M., Kox, T., Abbadi, A., Leckband, G., Snowdon, R. J., & Stahl, A. (2020). How Population Structure Impacts Genomic Selection Accuracy in Cross-Validation: Implications for Practical Breeding. Frontiers in Plant Science, 11(December), 1–14. https://doi.org/10.3389/fpls.2020.592977

Wimmer, V., Albrecht, T., Auinger, H. J., & Schön, C. C. (2012). Synbreed: A framework for the analysis of genomic prediction data using R. Bioinformatics, 28(15), 2086–2087. https://doi.org/10.1093/bioinformatics/bts335

Zheng, X., Levine, D., Shen, J., Gogarten, S. M., Laurie, C., & Weir, B. S. (2012). A high-performance computing toolset for relatedness and principal component analysis of SNP data. Bioinformatics, 28(24), 3326–3328. https://doi.org/10.1093/bioinformatics/bts606

