## Supplementary Table 1 for "Enviromic-based Kernels Optimize Resource Allocation with Multi-trait Multi-environment Genomic Prediction for Tropical Maize"

SUPPLEMENTARY MATERIAL

**Supplementary Table 1.** Prediction ability of the HEL dataset for single trait multi-environment trial (STMET) analysis for five models: EAD (M1); EAD+GE (M2); EADW (M3); EADW + GE (M4) and EADW + GW (M5), for both cross-validations CV1 and CV2. PAs were calculated by environment for grain yield (GY), ear height (EH) and plant height (PH). Values of PA are the average between environments. Standard deviation appears before the "±" sign.

| **Trait** |  | **M1** | **M2** | **M3** | **M4** | **M5** |
| --- | --- | --- | --- | --- | --- | --- |
| CV1 | | | | | | |
| GY | $\boldsymbol{\mu}$ | 0.53±0.20 | **0.66±0.13** | 0.53±0.20 | **0.66±0.13** | 0.64±0.15 |
|  | **%** | $-$20 | 0 | $-$20 | 0 | $-$3 |
| EH | $\boldsymbol{\mu}$ | 0.69±0.06 | 0.70±0.06 | 0.69±0.06 | **0.70±0.06** | 0.70±0.06 |
|  | **%** | $-$1 | - | $-$1 | **-** | - |
| PH | $\boldsymbol{\mu}$ | 0.72±0.06 | **0.73±0.06** | 0.72±0.06 | 0.73±0.06 | 0.73±0.06 |
|  | **%** | $-$2 | - | $-$2 | - | - |
| CV2 | | | | | | |
| GY | $\boldsymbol{\mu}$ | 0.52±0.17 | **0.68±0.12** | 0.52±0.18 | **0.68±0.12** | 0.67±0.13 |
|  | **%** | $-$24 | - | $-$24 | - | $-$2 |
| EH | $\boldsymbol{\mu}$ | 0.75±0.06 | 0.75±0.05 | 0.75±0.05 | **0.75±0.05** | 0.75±0.05 |
|  | **%** | - | - | - | **-** | - |
| PH | $\boldsymbol{\mu}$ | 0.78±0.05 | 0.79±0.05 | 0.78±0.05 | **0.79±0.05** | 0.79±0.05 |
|  | **%** | $-$ 2 | - | $-$2 | - | - |

**Supplementary Table 2.** Prediction ability of the HEL dataset for multi-trait multi-environment trial (MTMET) analysis for the five models under study: EAD (M1); EAD+GE (M2); EADW (M3); EADW + GE (M4) and EADW + GW (M5), for both cross-validation schemes CV1 and CV2. PAs were calculated by environment for grain yield (GY), ear height (EH) and plant height (PH). Values of PA are the average between environments. Standard deviation appears before the "±" sign.

| **Trait** |  | **M1** | **M2** | **M3** | **M4** | **M5** |
| --- | --- | --- | --- | --- | --- | --- |
| **CV1** | | | | | | |
| GY | $\boldsymbol{\mu}$ | 0.54±0.18 | 0.66±0.12 | 0.54±0.17 | **0.67±0.12** | 0.65±0.14 |
|  | **%** | $-$19 | - | $-$19 | - | $-$3 |
| EH | $\boldsymbol{\mu}$ | 0.70±0.06 | 0.70±0.06 | 0.70±0.06 | **0.71±0.06** | 0.70±0.06 |
|  | **%** | $-$1 | - | $-$1 | **-** | - |
| PH | $\boldsymbol{\mu}$ | 0.72±0.06 | 0.73±0.06 | 0.72±0.06 | **0.73±0.06** | 0.73±0.06 |
|  | **%** | $-$1 | - | $-$1 | - | - |
| **CV2** | | | | | | |
| GY | $\boldsymbol{\mu}$ | 0.53±0.16 | 0.68±0.11 | 0.53±0.16 | **0.68±0.11** | 0.67±0.12 |
|  | **%** | $-$22 | - | $-$22 | - | $-$1 |
| EH | $\boldsymbol{\mu}$ | 0.70±0.06 | 0.75±0.06 | 0.75±0.06 | **0.75±0.06** | 0.75±0.06 |
|  | **%** | - | - | - | **-** | - |
| PH | $\boldsymbol{\mu}$ | 0.78±0.05 | **0.79±0.05** | 0.78±0.05 | **0.79±0.05** | 0.79±0.05 |
|  | **%** | $-$1 | - | $-$1 | - | - |

Supplementary Table 3. Prediction ability of the USP dataset for single trait multi environment trial (STMET) analysis for the five models under study: EAD (M1); EAD+GE (M2); EADW (M3); EADW + GE (M4) and EADW + GW (M5) ), for both cross-validation schemes CV1 and CV2. PAs were calculated by environment for grain yield (GY), ear height (EH) and plant height (PH). Values of PA are the average between environments. Standard deviation appears before the "±" sign.

| **Trait** |  | **M1** | **M2** | **M3** | **M4** | **M5** |
| --- | --- | --- | --- | --- | --- | --- |
| **CV1** | | | | | | |
| GY | $\boldsymbol{\mu}$ | 0.46±0.06 | 0.48±0.05 | 0.46±0.06 | **0.48±0.05** | 0.47±0.05 |
|  | **%** | $-$4 | - | $-$4 | - | $-$1 |
| EH | $\boldsymbol{\mu}$ | 0.57±0.30 | **0.62±0.22** | 0.57±0.30 | **0.62±0.22** | 0.62±0.22 |
|  | **%** | $-$8 | **-** | $-$8 | **-** | - |
| PH | $\boldsymbol{\mu}$ | 0.65±0.05 | 0.65±0.05 | 0.65±0.05 | **0.65±0.05** | **0.65±0.05** |
|  | **%** | - | - | - | **-** | **-** |
| **CV2** | | | | | | |
| GY | $\boldsymbol{\mu}$ | 0.48±0.05 | **0.51±0.05** | 0.48±0.05 | **0.51±0.05** | 0.50±0.05 |
|  | **%** | $-$6 | **-** | $-$6 | **-** | $-$2 |
| EH | $\boldsymbol{\mu}$ | 0.58±0.31 | **0.61±0.24** | 0.58±0.31 | **0.61±0.24** | 0.61±0.25 |
|  | **%** | $-$5 | **-** | $-$5 | **-** | $-$1 |
| PH | $\boldsymbol{\mu}$ | 0.70±0.04 | 0.70±0.04 | 0.70±0.04 | 0.70±0.04 | **0.70±0.04** |
|  | **%** | - | - | - | - | **-** |

**Supplementary Table 4.** Prediction ability of the USP dataset for multi-trait multi environment trials (MTMET) analysis for the five models under study: EAD (M1); EAD+GE (M2); EADW (M3); EADW + GE (M4) and EADW + GW (M5) ), for both cross-validation schemes CV1 and CV2. PAs were calculated by environment for grain yield (GY), ear height (EH) and plant height (PH). Values of PA are the average between environments. Standard deviation appears before the "±" sign.

| **Trait** |  | **M1** | **M2** | **M3** | **M4** | **M5** |
| --- | --- | --- | --- | --- | --- | --- |
| **CV1** | | | | | | |
| GY | $\boldsymbol{\mu}$ | 0.46±0.06 | 0.47±0.05 | 0.46±0.06 | **0.48±0.05** | 0.47±0.05 |
|  | **%** | $-$4 | - | $-$4 | **-** | $-$2 |
| EH | $\boldsymbol{\mu}$ | 0.57±0.30 | 0.62±0.22 | 0.57±0.30 | **0.62±0.22** | 0.62±0.22 |
|  | **%** | $-$8 | - | $-$8 | **-** | - |
| PH | $\boldsymbol{\mu}$ | 0.65±0.05 | 0.65±0.05 | 0.65±0.05 | 0.65±0.05 | **0.65±0.05** |
|  | **%** | - | - | - | - | **-** |
| **CV2** | | | | | | |
| GY | $\boldsymbol{\mu}$ | 0.48±0.05 | 0.51±0.05 | 0.48±0.05 | **0.51±0.05** | 0.51±0.05 |
|  | **%** | $-$6 | - | $-$6 | **-** | - |
| EH | $\boldsymbol{\mu}$ | 0.58±0.31 | 0.61±0.25 | 0.58±0.31 | **0.61±0.25** | 0.61±0.25 |
|  | **%** | $-$5 | - | $-$5 | **-** | - |
| PH | $\boldsymbol{\mu}$ | 0.70±0.04 | 0.70±0.04 | 0.70±0.04 | 0.70±0.04 | **0.70±0.04** |
|  | **%** | - | - | - | - | **-** |
